# Dopamine release in nucleus accumbens is under tonic inhibition by adenosine A_1_ receptors regulated by astrocytic ENT1 and dysregulated by ethanol

**DOI:** 10.1101/2021.11.27.470186

**Authors:** Bradley M. Roberts, Elizabeth Lambert, Jessica A. Livesey, Zhaofa Wu, Yulong Li, Stephanie J. Cragg

## Abstract

Striatal adenosine A_1_ receptor (A_1_R) activation can inhibit dopamine release. A_1_Rs on other striatal neurons are activated by an adenosine tone that is limited by equilibrative nucleoside transporter 1 (ENT1) that is enriched on astrocytes and is ethanol-sensitive. We explored whether dopamine release in nucleus accumbens core is under tonic inhibition by A_1_Rs, and is regulated by astrocytic ENT1 and ethanol. In *ex vivo* striatal slices from male and female mice, A_1_R agonists inhibited dopamine release evoked electrically or optogenetically and detected using fast-scan cyclic voltammetry, most strongly for lower stimulation frequencies and pulse numbers, thereby enhancing the activity-dependent contrast of dopamine release. Conversely, A_1_R antagonists reduced activity-dependent contrast but enhanced evoked dopamine release levels, even for single optogenetic pulses indicating an underlying tonic inhibition. The ENT1 inhibitor NBTI reduced dopamine release and promoted A_1_R-mediated inhibition, and conversely, virally-mediated astrocytic overexpression of ENT1 enhanced dopamine release and relieved A_1_R-mediated inhibition. By imaging the genetically encoded fluorescent adenosine sensor GRAB-Ado, we identified a striatal extracellular adenosine tone that was elevated by the ENT1 inhibitor and sensitive to gliotoxin fluorocitrate. Finally, we identified that ethanol (50 mM) promoted A_1_R-mediated inhibition of dopamine release, through diminishing adenosine uptake via ENT1. Together, these data reveal that dopamine output dynamics are gated by a striatal adenosine tone, limiting amplitude but promoting contrast, regulated by ENT1, and promoted by ethanol. These data add to the diverse mechanisms through which ethanol modulates striatal dopamine, and to emerging datasets supporting astrocytic transporters as important regulators of striatal function.

**SIGNIFICANCE STATEMENT:** Dopamine axons in the mammalian striatum are emerging as strategic sites where neuromodulators can powerfully influence dopamine output in health and disease. We found that ambient levels of the neuromodulator adenosine tonically inhibit dopamine release in nucleus accumbens core via adenosine A_1_ receptors (A_1_Rs), to a variable level that promotes the contrast in dopamine signals released by different frequencies of activity. We reveal that the equilibrative nucleoside transporter 1 (ENT1) on astrocytes limits this tonic inhibition, and that ethanol promotes it by diminishing adenosine uptake via ENT1. These findings support the hypotheses that A_1_Rs on dopamine axons inhibit DA release and, furthermore, that astrocytes perform important roles in setting the level of striatal dopamine output, in health and disease.

## INTRODUCTION

Striatal dopamine (DA) axons are major strategic sites for striatal neuromodulators to influence DA output (Nolan et al., 2020; Rice et al., 2011; Roberts et al., 2021; Sulzer et al., 2016). Adenosine acts at A_1_- and A_2A_-receptors on diverse neurons in striatum, and exogenous activation of striatal A_1_-receptors (A_1_Rs) but not A_2_-receptors inhibits evoked DA release (Okada et al., 1996; O’Neill et al., 2007; O’Connor and O’Neill, 2008; Ross and Venton, 2015). Immunocytochemical studies of rat synaptosomes suggest that A_1_Rs can be localised directly to striatal DA axons (Borycz et al., 2007), but definitive evidence is lacking and other intermediary inputs to DA axons have not been excluded. In nucleus accumbens core (NAcC), adenosine can provide tonic A_1_R-mediated inhibition of glutamatergic and GABAergic transmission (Brundege and Williams, 2002; Adhikary and Birdsong, 2021). DA release in NAcC and caudate-putamen has been suggested to be under tonic A_1_R-mediated inhibition, as A_1_R antagonists increase extracellular DA levels in rats *in vivo* measured by microdialysis (Okada et al., 1996; Solinas et al., 2002; Quarta et al., 2004a, 2004b; Borycz et al., 2007), but whether these effects were direct and local, or involved intact long-loop circuits was not resolved. Furthermore, adenosine acts in a frequency-dependent manner on glutamate and GABA transmission across other nuclei, resulting in a stronger A_1_R-dependent inhibition of neurotransmission elicited by low-frequency versus high-frequency electrical stimulations (e.g. neocortex: Perrier et al., 2019; Qi et al., 2017; Yang et al., 2007; hippocampus: Moore et al., 2003; calyx of Held: Wong et al., 2006). It has not yet been established whether striatal A_1_Rs simply inhibit DA neurotransmission, or might also promote contrast in DA signals released by different firing rates, and whether in turn this is independent from striatal acetylcholine or GABA circuits which modify DA signal contrast (Rice and Cragg, 2004; Lopes et al., 2019).

Extracellular adenosine concentrations are limited by the activity of nucleoside transporters, most notably the equilibrative nucleoside transporter type 1 (ENT1) (Young et al., 2008; Nguyen et al., 2015). ENT1 is expressed in striatum (Jennings et al., 2001; Anderson et al., 2002) and is especially abundant on astrocytes (Peng et al., 2005; Chai et al., 2017). ENT1 on astrocytes can regulate adenosine signalling in striatum and elsewhere (Nagai et al., 2005; Tanaka et al., 2011; Boddum et al., 2016; Cheffer et al., 2018; Hong et al., 2020), and in dorsal striatum, astrocytic ENT1 activity modulates reward-seeking behaviours (Hong et al., 2020; Kang et al., 2020). We recently identified that astrocytes regulate DA release, owing to their expression of GABA transporters that regulate tonic GABAergic inhibition (Roberts et al., 2020), but whether ENT1 on astrocytes also modulates DA output by regulating adenosine tone at A_1_Rs has not been explored. Additionally, adenosine uptake by ENT1 is impaired by acute ethanol, which augments extracellular adenosine levels (Nagy et al., 1989, 1990; Choi et al., 2004) and contributes to the ataxic and hypnotic effects of ethanol through activation of striatal A_1_Rs (Meng and Dar, 1995; Phan et al., 1997; Dar, 2001). Acute ethanol reduces DA release evoked in NAcC in brain slices (Yorgason et al., 2014, 2015), as does chronic intermittent ethanol exposure in mice (Karkhanis et al., 2015; Rose et al., 2016), raising the question of whether ethanol regulates DA by promoting A_1_R-mediated inhibition of DA release.

Here, we assessed A_1_R regulation of DA release in NAcC by detecting DA in real-time using fast-scan cyclic voltammetry and detecting striatal adenosine using the genetically encoded fluorescent adenosine sensor (GRAB-Ado) (Peng et al., 2020; Wu et al., 2020). We reveal that A_1_Rs can tonically inhibit DA output though an ambient adenosine tone, and that A_1_Rs additionally regulate the activity-sensitivity of DA release, independently from striatal acetylcholine or GABA inputs. Furthermore, we find that tonic A_1_R-mediated inhibition of DA release is regulated by adenosine uptake by ENT1 on astrocytes, and dysregulated by ethanol.

## MATERIALS & METHODS

### Animals

All procedures were performed in accordance with the Animals in Scientific Procedures Act 1986 (Amended 2012) with ethical approval from the University of Oxford, and under authority of a Project Licence granted by the UK Home Office. Experiments were carried out using male and female adult (6-12 week-old) C57BL/6J mice (Charles River) or heterozygote knock-in mice bearing an internal ribosome entry site (IRES)-linked Cre recombinase gene downstream of the gene *Slc6a3*, which encodes the plasma membrane DA transporter (*Slc6a3^IRES-Cre^* mice; *B6.SJL-Slc6a3*^tm:L1(cre)Bkmn^/J; Jackson Laboratories stock no. 006660) maintained on a C57BL/6J background. All mice were group-housed and maintained on a 12-hr light cycle (light ON from 07:00 – 19:00) with *ad libitum* access to food and water. Data from male and female mice were combined throughout as no differences in the effects of adenosine A_1_ receptor agonists or antagonists between sexes were observed (**Table 1**).

**Table 1.** Comparison of effects of adenosine A_1_ receptor agonists and antagonists on evoked mean peak [DA]_o_ between sexes.

| Sex | Mean [DA]<sub>o</sub> (% pre-drug) | SEM | n | unpaired t test |
| --- | --- | --- | --- | --- |
| Figure 1B: A<sub>1</sub> agonist (CPA), electrically-evoked DA release |  |  |  |  |
| Male | 39.6 | 6.8 | 3 | $t_{(4)} = 0.467, p = 0.665$ |
| Female | 43.2 | 3.9 | 3 |  |
| Figure 1G: A<sub>1</sub> agonist (CPA), optogenetically-evoked DA release |  |  |  |  |
| Male | 59.4 | 2.9 | 4 | $t_{(6)} = 1.141, p = 0.297$ |
| Female | 53.4 | 4.4 | 4 |  |
| Figure 3A: A<sub>1</sub> antagonist (CPT), electrically-evoked DA release |  |  |  |  |
| Male | 140.8 | 7.8 | 4 | $t_{(5)} = 0.181, p = 0.864$ |
| Female | 142.6 | 5.4 | 3 |  |
| Figure 3C: A<sub>1</sub> antagonist (Caffeine), electrically-evoked DA release |  |  |  |  |
| Male | 117.4 | 8.0 | 3 | $t_{(4)} = 0.304, p = 0.777$ |
| Female | 114.9 | 2.2 | 3 |  |
| Figure 3E: A<sub>1</sub> antagonist (CPT), optogenetically-evoked DA release |  |  |  |  |
| Male | 126.6 | 16.7 | 3 | $t_{(6)} = 0.258, p = 0.805$ |
| Female | 130.0 | 3.8 | 5 |  |

### Stereotaxic intracranial injections

Mice were anesthetized with isoflurane and placed in a small animal stereotaxic frame (David Kopf Instruments). After exposing the skull under aseptic techniques, a small burr hole was drilled and adeno-associated viral solutions were injected at an infusion rate of 100 nL/min with a 32-gauge Hamilton syringe (Hamilton Company) using a microsyringe pump (World Precision Instruments) and withdrawn 5 min after the end of injection. To virally express ChR2 selectively in DA neurons, 600 nL per hemisphere of AAV5-EF1α-DIO-hChR2(H134R)-eYFP (8 × 10^12^ genome copies per ml; UNC Vector Core Facility) encoding Cre-dependent ChR2 was injected bilaterally into the midbrain (AP −3.1 mm, ML ± 1.2 mm from bregma, DV −4.25 mm from exposed dura mater) of 6 week-old *Slc6a3^IRES-Cre^* mice, following previously described methods (Roberts et al., 2020). To overexpress ENT1 in striatal astrocytes, 600 nL per hemisphere of AAV5-GfaABC_1_D-mENT1/mCherry-WPRE (1.6 x 10^13^ genome copies per ml; Vector Biolabs) encoding a fluorescence protein-fused and functional ENT1 under the astrocyte-specific abbreviated glial fibrillary acidic protein promoter (*GfaABC_1_D*) (Hong et al., 2020; Jia et al., 2020) or AAV5-GfaABC_1_D-mCherry-WPRE (6.1 x 10^12^ genome copies per ml; ETH Zurich Viral Vector Facility) for control fluorophore expression, was injected bilaterally into nucleus accumbens (AP +1.3 mm, ML ± 1.2 mm from bregma, DV −3.75 mm from exposed dura mater) of 6 week-old C57BL/6J wild-type mice. To virally express the fluorescent adenosine reporter GRAB-Ado (Peng et al., 2020; Wu et al., 2020), 800 nL per hemisphere of AAV9-hSyn-GRAB-Ado1.0m (0.5 x 10^13^ genome copies per ml; WZ Biosciences) was injected bilaterally into nucleus accumbens or dorsal striatum (AP +0.8 mm, ML ± 1.75 mm from bregma, DV −2.40 mm from exposed dura mater) of 6 week-old C57BL/6J wild-type mice. Mice were used for experiments 2-4 weeks post-intracranial injection.

### Brain slice preparation

Acute brain slices were obtained from 8 – 12-week-old mice using standard techniques. Mice were culled by cervical dislocation within 1 – 2 hrs after start of light ON period of light cycle and brains were dissected out and submerged in ice-cold cutting solution containing (in mM): 194 sucrose, 30 NaCl, 4.5 KCl, 1 MgCl_2_, 26 NaHCO_3_, 1.2 NaH_2_PO_4_, and 10 D-glucose. Coronal slices 300 μm-thick containing striatum were prepared from dissected brain tissue using a vibratome (VT1200S, Leica Microsystems) and transferred to a holding chamber containing artificial cerebrospinal fluid (aCSF) containing (in mM): 130 NaCl, 2.5 KCl, 26 NaHCO_3_, 2.5 CaCl_2_, 2 MgCl_2_, 1.25 NaH_2_PO_4_ and 10 glucose. Sections were incubated at 34 °C for 15 min before they were stored at room temperature (20–22 °C) until recordings were performed. All recordings were obtained within 8 h of slicing. All solutions were saturated with 95% O_2_/5% CO_2_. Before recording, individual slices were hemisected and transferred to a recording chamber and superfused at ~2.5–3.0 mL/min with aCSF at 31–33 °C.

### Fast-scan cyclic voltammetry (FSCV)

Evoked extracellular DA concentration ([DA]_o_) was measured in acute coronal brain slices using FSCV at carbon-fibre microelectrodes (7–10 μm diameter) fabricated in-house (tip length 70–120 μm) as used previously (Roberts et al., 2020). In brief, a triangular voltage waveform was scanned across the microelectrode (−700 to +1300 mV vs Ag/AgCl reference) at 800 V/s and at a scan frequency of 8 Hz using a Millar Voltammeter (Julian Millar, Barts and the London School of Medicine and Dentistry). Microelectrodes were calibrated post-hoc in 2 μM DA in each experimental solution. Microelectrode sensitivity to DA was between 10 and 40 nA/μM. Signals were attributed to DA due to the potentials of their characteristic oxidation (500-600 mV) and reduction (−200 mV) peaks. Currents at the oxidation peak potential were measured from the baseline of each voltammogram and plotted against time to provide profiles of [DA]_o_ versus time. Recordings were carried out in the nucleus accumbens core (NAcC), within ~100 μm of the anterior commissure, one site per slice. Electrical or light stimuli were delivered at 2.5 min intervals, which allow stable DA release to be sustained at ~90–95% of original levels over the typical time course of experiments, in control conditions (Roberts et al., 2020). In experiments where [DA]_o_ was evoked by electrical stimulation, a local bipolar concentric Pt/Ir electrode (25 μm inner diameter, 125 μm outer diameter; FHC Inc.) was placed ~100 μm from the recording microelectrode and stimulus pulses (200 μs duration) were given at 0.6 mA. We applied either single pulses (1p) or trains of 2–20 pulses at 10–100 Hz. In experiments where [DA]_o_ was evoked by light stimulation in slices prepared from *Slc6a3^IRES-Cre^* mice expressing ChR2, DA axons in striatum were activated by TTL-driven (Multi Channel Stimulus II, Multi Channel Systems) brief pulses (2 ms) of blue light (470 nm; 5 mWmm^-2^; OptoLED, Cairn Research), which illuminated the field of view (2.2 mm diameter, ×10 water-immersion objective). Data were digitized at 50 kHz using a Digidata 1550A digitizer (Molecular Devices). Data were acquired and analysed using Axoscope 11.0 (Molecular Devices) and locally written VBA scripts in Microsoft Excel (2013).

### GRAB-Ado Imaging

An Olympus BX51WI microscope equipped with a 470 nm OptoLED light system (Cairn Research), Iris 9 Scientific CMOS camera (Teledyne Photometrics), 525/50 nm emission filter (Cairn Research), and x10/0.3 NA water-immersion objective (Olympus) was used for wide-field fluorescence imaging of GRAB-Ado in striatal slices. Images acquisition was controlled using Micro-Manager 1.4. Electrical stimulations, LED light, and image acquisition were synchronised using TTL-driven stimuli via Multi Channel Stimulus II (Multi Channel Systems). Image files were analysed with Matlab R2017b and Fiji 1.5. For experiments measuring changes to basal, non-stimulated extracellular adenosine levels, images (100 ms exposure duration) were acquired every 30 s during brief exposure (1 s) to blue light (470 nm; 1 mWmm^-2^; OptoLED, Cairn Research) for a 40 min window. We extracted fluorescence intensity from the region of interest (ROI; 150 x 150 μm) and derived a background-subtracted fluorescence intensity (F_t_) by subtracting background fluorescence intensity from an equal-sized ROI where there was no GRAB-Ado expression (i.e. cortex). Data are expressed as a change in fluorescence (ΔF/F_0_) and were derived by calculating [(F_t_-F_0_)/F_0_], where F_0_ is the average background-subtracted fluorescence intensity (F_t_) of the first 20 acquired images (initial 10 mins). For experiments measuring extracellular adenosine levels in response to trains of electrical stimulation or experiments calibrating GRAB-Ado signals to applications of known concentrations of exogenous adenosine, images were acquired at 10 Hz (100 ms exposure duration) during continuous blue light (470 nm; 1 mWmm^-2^; OptoLED, Cairn Research) for a 4 min recording window. For evoked adenosine release, electrical stimulus pulses (100 pulses at 50 Hz, 200 μs pulse duration, 0.6 mA) were given by a local bipolar concentric Pt/Ir electrode (25 μm inner diameter, 125 μm outer diameter; FHC Inc.) at the 30 second timepoint. Bath applications of exogenous adenosine were also applied at the 30 second timepoint. Background-subtracted fluorescence intensity (F_t)_ was extracted from an ROI (150 x 150 μm) ~50 μm from the stimulating electrode and ΔF/F_0_ was derived by calculating [(F_t_-F_0_)/F_0_], where F_0_ is the average fluorescence intensity over the 10 s window (100 images) prior to onset of electrical stimulation or bath application of exogenous adenosine.

### Drugs

Adenosine (Ado, 25-100 μM), (+)-bicuculline (10 μM), dihydro-β-erythroidine hydrobromide (DHβE, 1 μM), nitrobenzylthioinosine (NBTI, 10 μM), and tetrodotoxin (TTX, 1 μM) were obtained from Tocris Bioscience. CGP 55845 hydrochloride (CGP, 4 μM), 8-Cyclopentyl-1,3-dimethylxanthine (CPT, 10 μM), and dipropylcyclopentylxanthine (DPCPX, 2 μM) were obtained from Abcam. Ethanol (EtOH, 50 mM), and N^6^-cyclopentyladenosine (CPA, 15 μM) were obtained from Sigma-Aldrich. Fluorocitrate (FC) was prepared as previously described (Paulsen et al., 1987; Roberts et al., 2020). In brief, D,L-fluorocitric acid Ba_3_ salt (Sigma–Aldrich) was dissolved in 0.1 M HCl, the Ba^2+^ precipitated with 0.1 M Na_2_S0_4_ and then centrifuged at 1000 × g for 5 min. Supernatant containing fluorocitrate was used at a final concentration of 100 μM for experimentation.

### Immunocytochemistry

To confirm viral ENT1-mCherry expression in non-neuronal cells, direct mCherry fluorophore expression was compared to indirect immunofluorescence for neuronal marker NeuN. ENT1-mCherry expressing mice were anaesthetised with an overdose of pentobarbital and transcardially perfused with phosphate-buffered saline (PBS), followed by 4% paraformaldehyde in 0.1 M phosphate buffer, pH 7.4. Brains were removed and post-fixed overnight in 4% PFA. Coronal sections were cut using a vibrating microtome (Leica VT1000S) at a thickness of 50 μm and collected in a 1 in 4 series. Sections were stored in PBS with 0.05% sodium azide until processing. Upon processing, sections were washed in PBS and then blocked for 1 h in a solution of PBS TritonX (0.3%; PBS-Tx) containing 10% normal donkey serum (NDS). Sections were then incubated in primary antibodies overnight in PBS-Tx with 2% NDS at 4 °C. Primary antibodies: rabbit anti-NeuN (1:500, Biosensis, R-3770–100). Sections were then incubated in species-appropriate fluorescent secondary antibodies with minimal cross-reactivity for 2 hours in PBS-Tx with 2% NDS at room temperature. Secondary antibodies: Donkey anti-rabbit AlexaFluor 405 (1:500, Abcam, ab175651). Sections were washed in PBS and then mounted on glass slides and cover-slipped using Vectashield (Vector Labs). Coverslips were sealed using nail varnish and stored at 4 °C. Confocal images were acquired with an Olympus FV3000 Confocal Laser Scanning Microscope using a x20 and x40 objective and filters for appropriate excitation and emission wave lengths (Olympus Medical).

## RESULTS

### Adenosine A_1_Rs inhibit striatal DA release

Previous studies have shown that agonists for striatal A_1_Rs inhibit DA release evoked by discrete electrical stimulations in dorsal and ventral striatum (O’Neill et al., 2007; O’Connor and O’Neill, 2008; Ross and Venton, 2015). We firstly corroborated these observations in NAcC for DA release evoked by single stimulus pulses and detected using FSCV in *ex vivo* coronal slices (**Figure 1A**). Bath application of A_1_R agonist CPA (15 μM) reduced evoked [DA]_o_ by ~50% (**Figure 1B**, t_(5)_ = 15.2, *p* < 0.0001, paired t test). This effect of CPA was prevented by prior application of A_1_R antagonist DPCPX (2 μM) (**Figure 1B**, F_(1, 11)_ = 88.6, *p* < 0.0001, two-way RM ANOVA for effect of CPA in the absence vs presence of DPCPX; **Figure 1E**, t_(11)_ = 9.99, *p* < 0.0001, unpaired t test), confirming an action via A_1_Rs.

**Figure 1.**
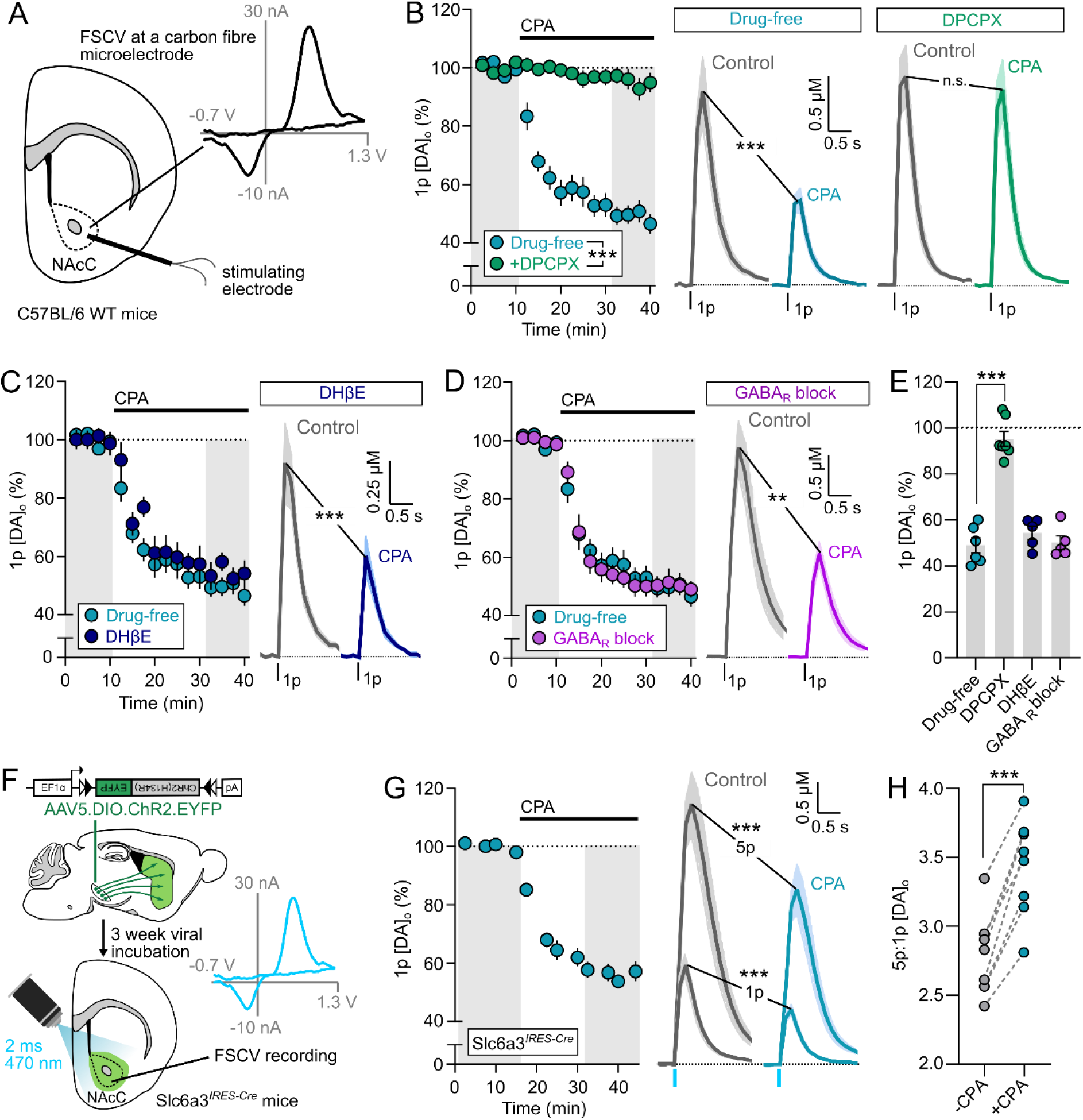
Adenosine A_1_R agonists inhibit DA release in NAcC. **A**, Experimental setup for FSCV for DA detection in wild-type mouse NAcC in acute slices. Inset, typical evoked DA voltammogram. **B-D**, Left, Summary of mean peak [DA]_o_ evoked by a single electrical pulse (1p) before and after application of A_1_R agonist CPA (15 μM) normalised to pre-drug baseline (dotted line). Shaded areas represent time points used to obtain data on right, mean [DA]_o_ transients in either drug-free ACSF (blue *n* = 6 experiments/4 mice), or in the presence of A_1_R antagonist DPCPX (2 μM) (B, green, *n* = 8 experiments/5 mice), or nAChR antagonist DHβE (1 μM) (C, dark blue, *n* = 5 experiments/4 mice), or GABAA and GABAB antagonists (+)-bicuculline (10 μM) and CGP 55845 (4 μM) (D, purple, *n* = 5 experiments/4 mice). Control data in (C) and (D) are from (B). **E**, Mean peak [DA]_o_ following application of CPA (15 μM) (as a % of pre-drug baseline, data summarised from (B-D)). **F**, Viral injection into midbrain of Slc6a3^*IRES-Cre*^ mice for expression of ChR2-eYFP in DA axons for optogenetic-evoked DA release. **G**, As in (B) but [DA]_o_ evoked optogenetically by single light pulses (1p) or five pluses of light (5p) at 25 Hz (*n* = 8 experiments/5 mice). **H**, Ratio of peak [DA]_o_ evoked by 5p:1p at 25 Hz before (grey) and after (blue) application of CPA (15 μM). Statistics: \*\*\**p*<0.001, paired t tests (B,C,D,G,H), unpaired t tests (E) and two-way repeated measures ANOVA (B,C,D). Error bars indicate SEM.

Striatal cholinergic interneurons (ChIs) have A_1_Rs (Alexander and Reddington, 1989; Ferré et al., 1996; Song et al., 2000; Preston et al., 2009) through which A_1_R agonists can hyperpolarise ChIs and inhibit ACh release (Richardson and Brown, 1987; Brown et al., 1990; Preston et al., 2009). Because ChIs operate strong control over DA release via nAChRs (Jones et al., 2001; Rice and Cragg, 2004; Zhang and Sulzer, 2004), and can mediate the effects of other striatal neuromodulators on DA (Britt and McGehee, 2008; Hartung et al., 2011; Stouffer et al., 2015; Kosillo et al., 2016; Lemos et al., 2019), they might indirectly mediate the control of DA by A_1_Rs. We explored whether A_1_Rs can modulate striatal DA release in the absence of these potential actions on ChIs and nAChRs. We used nAChR antagonist DHβE (1 μM) to inhibit nAChRs as described previously (Rice and Cragg, 2004; Threlfell et al., 2012), after which subsequent application of CPA nonetheless reduced evoked [DA]_o_ evoked by single stimulus pulses by ~50% (**Figure 1C**, t_(4)_ = 15.9, *p* < 0.0001, paired t test), an effect not different from that seen in the absence of DHβE (**Figure 1C**, F_(1, 9)_ = 1.22, *p* = 0.29, two-way RM ANOVA: main effect of drug; **Figure 1E**, t_(9)_ = 1.21, *p* = 0.259, unpaired t test), indicating that A_1_Rs can suppress DA release independently from any indirect effects via ChI inputs to nAChRs.

DA release throughout striatum is also under tonic inhibition by GABA through action at GABAA and GABAB receptors (Lopes et al., 2019; Roberts et al., 2020), and we tested whether A_1_Rs require this GABA input to inhibit DA release. However, in the presence of GABAA and GABAB receptor antagonists, bicuculline (10 μM) and CGP 55845 (5 μM) respectively, the A_1_R agonist CPA (15 μM) significantly reduced [DA]_o_ evoked in NAcC by single electrical pulses by ~40% (**Figure 1D**, t_(4)_ = 6.10, *p* = 0.0036, paired t test), an effect that was not significantly different from that seen in the absence of GABA receptor antagonists (**Figure 1D**, F_(1, 9)_ = 0.019, *p* = 0.892, two-way RM ANOVA: main effect of drug; **Figure 1E**, t_(9)_ = 0.259, *p* = 0.802, unpaired t test), indicating that A_1_ receptor agonists can inhibit DA release independently from indirect effects acting via striatal GABA modulation of DA.

To further test whether the inhibition of DA release by A_1_Rs requires the co-activation of other striatal neuron types or inputs, we tested whether A_1_Rs modulate DA release evoked optogenetically by targeted light activation of DA axons. Optogenetic activation of DA axons minimises the activation of other striatal neurons that occurs with non-selective electrical stimulation; optogenetically evoked DA release is independent of at least nAChR input (Threlfell et al., 2012; Melchior et al., 2015). We expressed ChR2-eYFP in DA neurons and axons in Slc6a3*^IRES-Cre^* mice using an established viral approach (**Figure 1F**) as we previously described (Roberts et al., 2020). CPA suppressed [DA]_o_ evoked in NAcC by single brief (2 ms) blue light pulses (**Figure 1G**, t_(7)_ = 16.2, *p* < 0.0001, paired t test), indicating that A_1_Rs can inhibit DA release independently from not just ChI input to nAChRs but also without requiring the coincident activation of inputs from other neuron types, suggesting that A_1_R inhibition of DA release could be via direct action on DA axons, although actions via an unidentified tonically active input that is not GABA are possible.

### Striatal A_1_Rs enhance the frequency- and activity-sensitivity of DA release

Previous studies exploring whether striatal A_1_Rs regulate DA release have predominantly focused on the effects on release evoked by single, discrete stimuli, and impact on release by the full range of physiological relevant frequencies has not been explored. Adenosine acts in a frequency-dependent manner on glutamate and GABA transmission in other brain nuclei, resulting in a stronger A_1_R-dependent inhibition of neurotransmission elicited by low-frequency versus high-frequency electrical stimulations, and therefore operating like a high-pass input filter on neurotransmitter release (e.g. neocortex: Perrier et al., 2019; Qi et al., 2017; Yang et al., 2007; hippocampus: Moore et al., 2003; calyx of Held: Wong et al., 2006). Other inputs and mechanisms that inhibit DA release by single stimuli can also promote the frequency dependence of DA release e.g. GABA (Lopes et al., 2019; Roberts et al., 2020) consistent with a reduction in the initial release probability and a consequent relief of some short-term depression (Condon et al., 2019). Using optogenetic activation, we observed a sizeable enhancement in the 5p:1p ratio (for 25 Hz) of light-evoked [DA]_o_ in the NAcC following application of CPA (**Figure 1H**, t_(7)_ = 10.4, *p* < 0.0001, paired t test), suggesting that striatal A_1_Rs might promote the sensitivity of DA release to firing rate.

We used electrical stimulation to investigate the effects of A_1_R activation on DA release across a full range of physiological DA neuron firing frequencies (1, 10, 50 and 100 Hz), with varying pulse numbers (5, 10, 15 and 20 p). These experiments were carried out in the presence of nAChR antagonist DHβE (1 μM) to eliminate the effects of ChIs which profoundly restrict the apparent frequency- and activity-sensitivity of evoked DA release (Rice and Cragg, 2004; Zhang and Sulzer, 2004). A_1_R activation by CPA (15 μM) inhibited DA release inversely and significantly with stimulation frequency (**Figure 2A-B**, F_(3, 18)_ = 53.9, *p* < 0.0001, one-way RM ANOVA), such that CPA significantly increased the frequency-dependence of evoked DA release (**Figure 2C**, F_(1, 7)_ = 38.2, *p* = 0.0005, two-way RM ANOVA: main effect of drug; F_(3, 21)_ = 23.0, *p* < 0.0001, frequency x drug interaction). In addition, A_1_R activation by CPA inhibited DA release in a manner that varied inversely with number of pulses in a train (50 Hz) (**Figure 2D-E**, F_(6, 24)_ = 31.1, *p* < 0.0001, one-way RM ANOVA) which significantly increased the pulse-dependence of evoked DA release (**Figure 2F**, F_(1, 7)_ = 46.7, *p* = 0.0002, two-way RM ANOVA: main effect of drug; F_(4, 28)_ = 28.2, *p* < 0.0001, pulse number x drug interaction). Therefore, striatal A_1_R do not simply inhibit DA release, but have a dynamic outcome depending on activity, that promotes the contrast in DA signals released by different activity.

**Figure 2.**
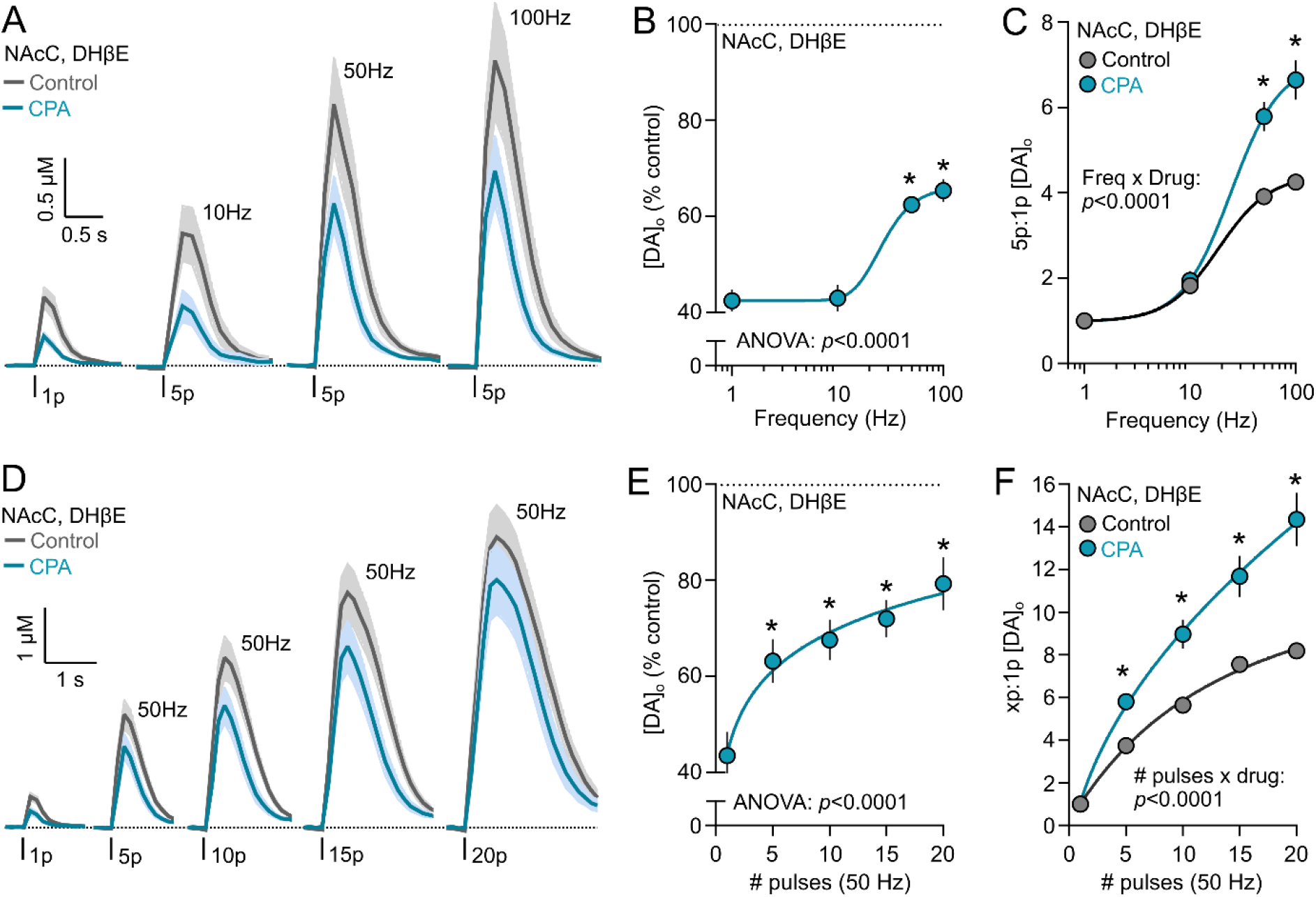
Adenosine A_1_R agonists modify the frequency- and activity-sensitivity of DA release in NAcC. **A**, Mean [DA]_o_ transients evoked by 1 or 5 electrical pulses in control conditions (grey) or in A_1_R agonist CPA (15 μM) (blue) in NAcC (*n* = 7 experiments/5 mice). **B**, Mean peak [DA]_o_ from (A) normalised to control conditions versus stimulation frequency. **C**, Mean peak [DA]_o_ from (A) normalised to 1p in each condition versus stimulation frequency. **D**, Mean [DA]_o_ transient evoked by 1-20 pulses (50 Hz) of electrical trains in control conditions (grey) or in A_1_R agonist CPA (15 μM) (blue) in NAcC (*n* = 7 experiments/5 mice). **E**, Mean peak [DA]_o_ from (D) normalised to control conditions versus pulse number. **F**, Mean peak [DA]_o_ from (D) normalised to 1p in each condition versus number of pulses. Statistics: \**p* < 0.001, one-way repeated measures ANOVA with Sidak’s multiple comparisons to 1p (B,E), two-way repeated measures ANOVA with Sidak’s multiple comparisons (C,F). Sigmoidal non-linear curve fits. Error bars indicate SEM. DHβE (1 μM) present throughout.

### Adenosine operates a tonic inhibition on DA release

We next addressed whether A_1_Rs provide an endogenous inhibition of DA release levels and support the contrast in DA signals released by different activity. Previous *in vivo* microdialysis studies indicate that endogenous adenosine might even exert a tonic A_1_R-mediated inhibition of DA release, as systemic dosing or intrastriatal infusions of A_1_R antagonists (as well as competitive A_1_ and A_2A_ receptor antagonist caffeine (Solinas et al., 2002; Quarta et al., 2004b)), increase extracellular striatal DA levels (Okada et al., 1996; Quarta et al., 2004a; Borycz et al., 2007). These effects might involve indirect actions via long loop circuits that are intact *in vivo* that could modulate DA neuron firing in midbrain. We tested therefore whether endogenous inhibition of DA release by A_1_Rs could be localised to the NAcC and whether it occurs without co-activation of other neurons, indicative of an tonic inhibition of DA release, by exploring the effects of A_1_R antagonists on DA release in NAcC evoked either electrically or optogenetically in coronal striatal slices.

Application of the A_1_R antagonist CPT (10 μM) significantly enhanced [DA]_o_ evoked by single or five pulses (50 Hz) of electrical stimulation by ~30-40% (**Figure 3A**, 1p: t_(6)_ = 5.10, *p* = 0.0022; 5p: t_(6)_ = 5.13, *p* = 0.0022; paired t tests), and in turn, decreased the ratio of [DA]_o_ evoked by 5p:1p (**Figure 3B**, t_(6)_ = 3.72, *p* = 0.0099, paired t test), indicative of an underlying regulation of DA release by endogenous adenosine. Caffeine (20 μM) also significantly enhanced [DA]_o_ evoked by single or five pulses (50 Hz) of electrical stimulation by ~20-10% (**Figure 3C**; 1p: t_(5)_ = 3.69, *p* = 0.014; 5p: t_(5)_ = 3.49, *p* = 0.018; paired t tests) and decreased the ratio of [DA]_o_ evoked by 5p:1p ratio (**Figure 3D**, t_(5)_ = 3.59, *p* = 0.016, paired t test). We then investigated whether A_1_R-mediated inhibition of DA release by endogenous adenosine might arise through an ambient adenosine tone, by exploring whether A_1_R antagonist CPT could promote DA release when evoked using optogenetic stimulation to activate DA axons selectively without co-activation of other striatal cell types that might provide a source of endogenous adenosine. CPT significantly enhanced [DA]_o_ evoked by single and five pulses (25 Hz) of light in Slc6a3^*IRES-Cre*^ ChR2-expressing mice by ~20-30% (**Figure 3E**, 1p: t_(7)_ = 5.58, *p* = 0.0008; 5p: t_(7)_ = 5.51, *p* = 0.0009; paired t tests) and decreased the ratio of [DA]_o_ evoked by 5p:1p (25 Hz) (**Figure 3F**, t_(7)_ = 13.50, *p* < 0.0001, paired t test). These data therefore suggest that striatal DA release is under tonic inhibition by an ambient adenosine tone at A_1_Rs, which promotes contrast in DA signals released by different activity.

**Figure 3.**
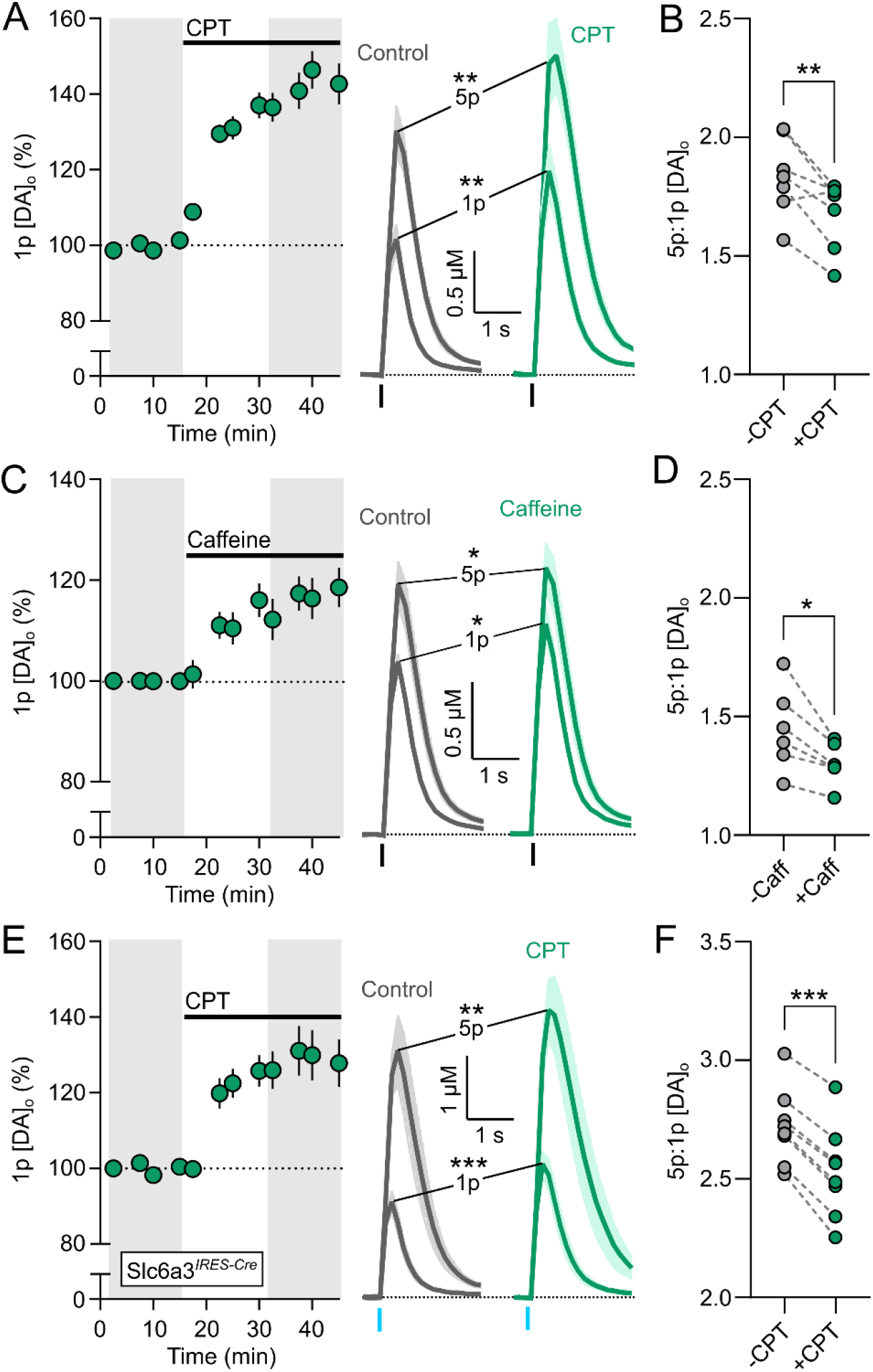
Adenosine A_1_R antagonists enhance DA release and correspondingly reduce the activity-sensitivity of DA release in NAcC. **A**, Left, summary of mean peak [DA]_o_ before (grey) and after application of A_1_R antagonist CPT (10 μM) (green) normalised to pre-drug baseline (dotted line) and right, corresponding mean [DA]_o_ transients evoked by a single electrical pulse (1p) and five pulses (5p) at 50 Hz in NAcC (*n* = 7 experiments/4 mice). Shaded areas are used to obtain illustrated transients of [DA]_o_ and for statistical comparisons. **B**, Ratio of peak [DA]_o_ evoked by 5p:1p at 50 Hz before (grey) and after (blue) application of CPT (10 μM). Data summarised from (A). **C-D**, As in (A-B) but before (grey) and after application caffeine (20 μM) (green) (*n* = 6 experiments/4 mice). **E-F**, As in (A-B) but [DA]_o_ evoked optogenetically by single light pulses (1p) or five pulses (5p) of light at 25 Hz in Slc6a3^IRES-Cre^ mice (*n* = 8 experiments/5 mice). Statistics: \**p*<0.05, \*\**p*<0.01, \*\*\**p*<0.001, Paired *t* tests. Error bars indicate SEM.

### ENT1 on astrocytes is a regulator of tonic A_1_R-mediated inhibition of DA release

We next tested the hypothesis that striatal ENT1 and, by association, astrocytes, by governing ambient adenosine levels (Nagai et al., 2005; Young et al., 2008; Tanaka et al., 2011; Nguyen et al., 2015; Cheffer et al., 2018; Hong et al., 2020), might determine the level of tonic A_1_R-mediated inhibition of evoked DA release in NAcC. We first inhibited ENT1 activity by pre-treatment of slices with the selective ENT1 inhibitor NBTI (10 μM, for 45 – 60 min), which has previously been shown to increase adenosine levels in striatum (Pajski and Venton, 2010). ENT1 inhibition itself attenuated [DA]_o_ evoked by single and five pulses (50 Hz) of electrical stimulation (**Figure 4A**; 1p: t_(12)_ = 4.38, *p* = 0.0009; 5p: t_(12)_ = 3.02, *p* = 0.0107; unpaired t tests). Furthermore, after pre-treatment with NBTI, the A_1_R antagonist CPT (10 μM) enhanced [DA]_o_ evoked by single electrical pulses to a significantly greater degree than in control slices (**Figure 4B**, F_(1, 12)_ = 27.17, *p =* 0.0002, two-way RM ANOVA: main effect of drug). CPT decreased the ratio of [DA]_o_ evoked by 5p:1p (50 Hz) in both conditions (**Figure 4C**, F_(1, 12)_ = 35.48, *p* < 0.0001, two-way RM ANOVA: main effect of drug), but there was a significant statistical interaction between NBTI and CPT (**Figure 4C**, F_(1, 12)_ = 9.48, *p* = 0.0096, two-way RM ANOVA: NBTI x CPT interaction) borne out by a greater decrease in this ratio in slices pre-treated with NBTI compared to control slices (**Figure 4D**, t_(12)_ = 3.08, *p* = 0.0096, unpaired t test). These data suggest that ENT1 regulates, and in particular limits, how A_1_ receptors tonically inhibit DA release and support its associated activity-sensitivity.

**Figure 4.**
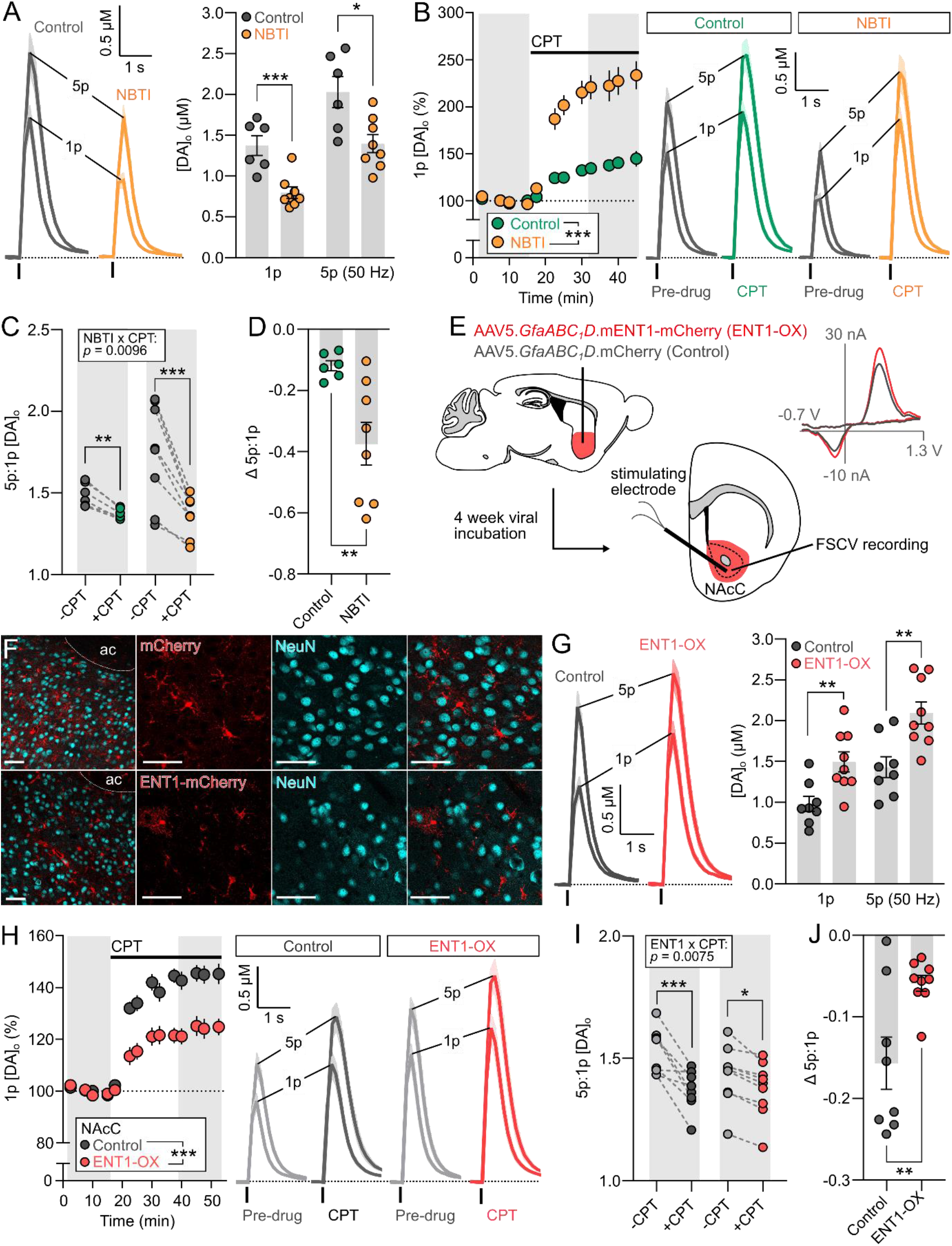
Adenosine uptake by ENT1 on striatal astrocytes regulates tonic A_1_R-mediated inhibition of DA release. **A**, Mean [DA]o transients (left) and corresponding mean peak [DA]o (right) evoked by a single electrical pulse (1p) or five pulses (5p) at 50 Hz in NAcC in the absence (grey; *n* = 6 experiments/5 mice) or presence of ENT1 inhibitor NBTI (10 μM) (orange; *n* = 8 experiments/5 mice). **B**, Summary of mean peak [DA]o before (grey) and after application of A_1_R antagonist CPT (10 μM) normalised to pre-drug baseline (dotted line) (left) and right, corresponding mean [DA]o transients evoked by a single electrical pulse (1p) and five pulses (5p) at 50 Hz in NAcC in the absence (green, *n* = 6 experiments/5 mice) or presence of ENT1 inhibitor NBTI (10 μM) (orange; *n* = 8 experiments/5 mice). **C**-**D**, Ratio of peak [DA]o evoked by 5p:1p at 50 Hz (C) and Δ 5p/1p (D) before (grey) and after application of CPT (10 μM) in absence (left, green) or presence of ENT1 inhibitor NBTI (10 μM) (right, orange). Data summarised from (B). **E**, Delivery to NAcC of viral fluorescence-tagged ENT1 driven by an astrocytic promoter to test impact of increased astrocyte-specific expression of ENT1 on DA release detected locally in NAcC. **F**, Immunofluorescence for mCherry (top, red), mCherry-tagged ENT1 (bottom, red) and neuronal marker NeuN (cyan) in NAcC. Scale bars: 40 μm. ac, anterior commissure. **G**, Mean [DA]_o_ transients (left) and corresponding mean peak [DA]_o_ (right) evoked by a single electrical pulse (1p) or five pulses (5p) at 50 Hz in NAcC in slices overexpressing ENT1 (ENT1-OX; red; *n* = 9 experiments/4 mice) or expressing control fluorophore mCherry (dark grey; *n* = 8 experiments/4 mice) under an astrocyte-specific promoter. **H**, Summary of mean peak [DA]_o_ before (grey) and after application of A_1_R antagonist CPT (10 μM) normalised to pre-drug baseline (dotted line) (left) and right, corresponding mean [DA]_o_ transients evoked by a single electrical pulse (1p) and five pulses (5p) at 50 Hz in NAcC in slices overexpressing ENT1 (ENT1-OX; red; *n* = 9 experiments/4 mice) or expressing control fluorophore mCherry (dark grey; *n* = 8 experiments/4 mice). **I**-**J**, Ratio of peak [DA]_o_ evoked by 5p:1p at 50 Hz (I) and Δ 5p:1p (J) before (light grey) and after application of CPT (10 μM) in slices overexpressing ENT1 (right, red) or expressing control fluorophore mCherry (left, dark grey). Data summarised from (H). Statistics: *p*<0.05, \*\**p*<0.01, \*\*\**p*<0.001, unpaired *t* test (A,D,G,J), two-way repeated measures ANOVA (B,C,H,I) with Sidak multiple comparisons (C,I). Error bars indicate SEM.

We then tested conversely whether an upregulation of ENT1 specifically on astrocytes could diminish the level of endogenous A_1_R-mediated inhibition of DA release. We targeted fluorescence-tagged ENT1 to astrocytes in the NAcC (**Figure 4E-F**) using a viral approach already validated for striatal astrocytes (Hong et al., 2020; Jia et al., 2020). Astrocyte-targeted ENT1-overexpression (ENT1-OX) increased [DA]_o_ evoked by single and five pulses (50 Hz) of electrical stimulation compared to targeted mCherry controls (**Figure 4G**, 1p: t_(15)_ = 3.29, *p* = 0.0049; 5p: t_(15)_ = 3.49, *p* = 0.0033; unpaired t tests). Furthermore, with ENT1-OX, the A_1_R antagonist CPT (10 μM) enhanced [DA]_o_ evoked by single electrical pulses to a significantly lesser degree than in mCherry controls (**Figure 4H**, F_(1, 15)_ = 20.12, *p* = 0.0004, two-way RM ANOVA: main effect of drug). CPT decreased the ratio of [DA]_o_ evoked by 5p:1p (50 Hz) in ENT1-OX and mCherry controls (**Figure 4I**, F_(1, 15)_ = 46.28, *p* < 0.0001, two-way RM ANOVA: main effect of drug), but there was a significant statistical interaction between ENT1-OX and CPT (**Figure 4I**, F_(1, 15)_ = 9.54, *p* = 0.0075, two-way RM ANOVA: ENT1-OX x CPT interaction), which was borne out by a smaller decrease in this ratio following CPT application in ENT1-OX than in controls (**Figure 4J**, t_(15)_ = 3.09, *p* = 0.0075, unpaired t test). Together, these data suggest that ENT1 on astrocytes in particular can support adenosine uptake and set the level of inhibition and regulation of activity-dependence of DA release by ambient adenosine acting at A_1_Rs.

To resolve directly whether ENT1, and astrocytes, govern striatal adenosine levels, we detected adenosine levels by imaging the recently developed GRAB-Ado sensor (Peng et al., 2020; Wu et al., 2020), a virally expressed genetic reporter, injected into striatum (**Figure 5A**). We confirmed that striatal GRAB-Ado sensor fluorescence responded to adenosine concentrations in a concentration-dependent manner on application of exogenous adenosine to slices (**Figure 5B**, F_(3,35)_ = 28.73, *p* < 0.0001, one-way ANOVA), and with a large range of dF/F. To assess the impact of ENT1 on ambient adenosine tone in NAcC, we imaged GRAB-Ado fluorescence before and after ENT1 inhibition, in the absence of any stimulation. Bath application of ENT1 inhibitor NBTI (10 μM) increased fluorescence compared to vehicle controls (**Figure 5C**, F_(1,18)_ = 57.8, *p* < 0.0001, two-way RM ANOVA: main effect of NBTI), indicating that ENT1 limits extracellular adenosine levels. To test whether ENT1 on astrocytes participate in this mechanism, we tested whether metabolic inhibition of astrocytes limited ENT1 function. We pre-treated slices with the gliotoxin fluorocitrate (100 μM, for 45 – 60 min), which has been established to induce metabolic arrest in astrocytes, render them inactive and prevent the effects of astrocytic transporters (Paulsen et al., 1987; Henneberger et al., 2010; Bonansco et al., 2011; Boddum et al., 2016; Roberts et al., 2020). The effects of ENT1 inhibitor NBTI on GRAB-Ado fluorescence were occluded after pre-treatment with fluorocitrate (**Figure 5D**, F_(1,14)_ = 14.32, *p* = 0.002, two-way RM ANOVA: main effect of FC).

**Figure 5.**
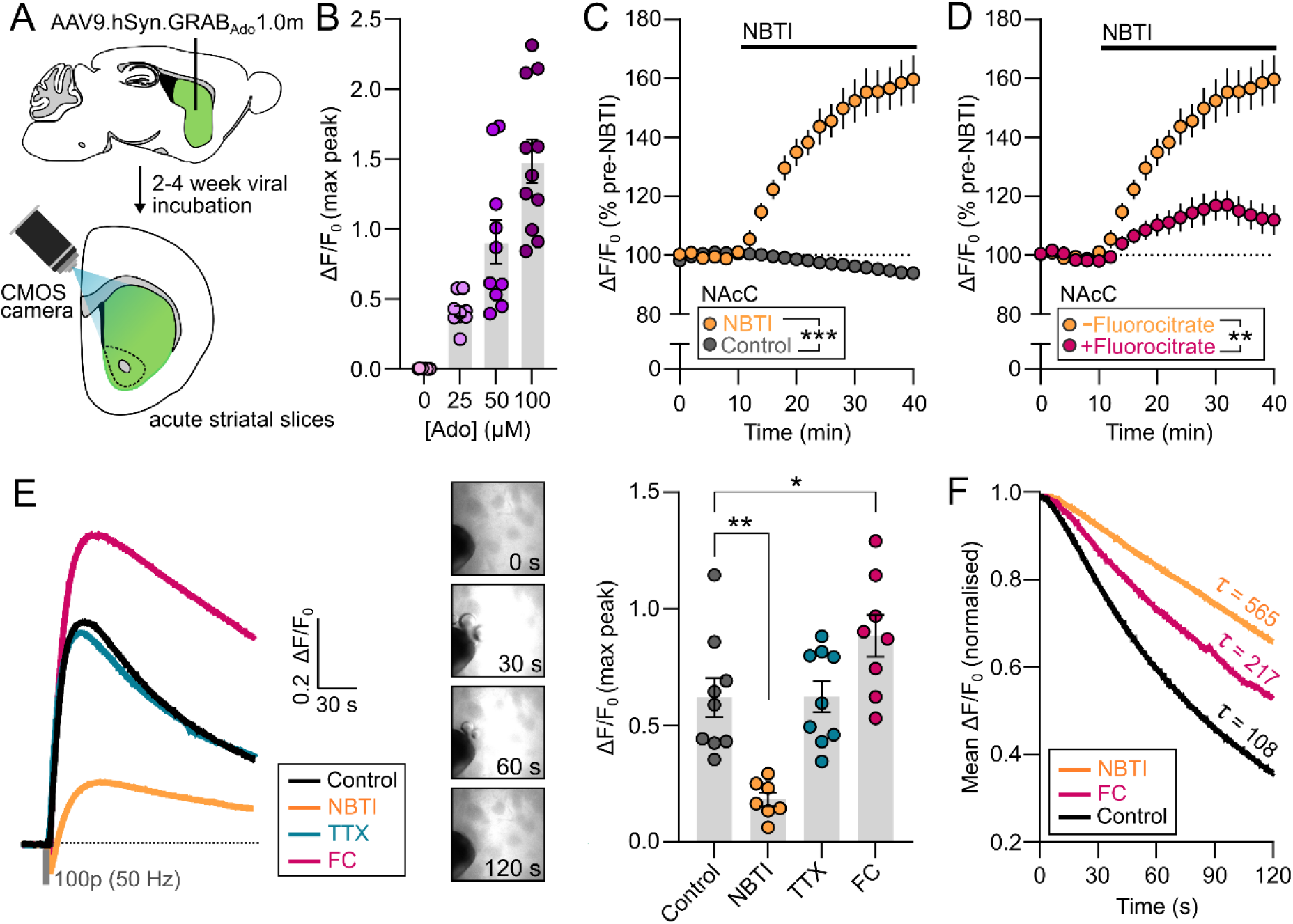
Striatal adenosine tone is regulated by ENT1 activity on striatal astrocytes. **A**, Viral delivery of GRAB-Ado to striatum for imaging extracellular adenosine levels *ex vivo* in acute striatal slices. **B**, Peak ΔF/F_0_ in response to increasing concentrations of exogenously applied adenosine (0 – 100 μM) (*n* = 9 – 11 slices/4 – 6 mice). **C**, Non-stimulated ΔF/F_0_ GRAB_Ado_ signal in NAcC normalised to before drug application (dotted line) in control conditions (dark grey, *n* = 10 experiments/5 mice) or before and after application of NBTI (10 μM) (orange, *n* = 10 experiments/5 mice). **D**, As in (C) but in the presence of gliotoxin fluorocitrate (100 μM) (pink, *n* = 6 experiments/ 4 mice). **E**, Mean transients (left) and mean peak (right) ΔF/F_0_ GRAB_Ado_ signal evoked by 100 electrical pulses (50 Hz) in drug-free control conditions (*n* = 9 slices / 5 mice) or the presence of NBTI (10 μM) (orange; *n* = 7 experiments/5 mice), TTX (1 μM) (blue; *n* = 9 experiments/5 mice) or fluorocitrate (100 μM) (pink; *n* = 8 experiments/5 mice). **F**, Mean ΔF/F_0_ decay phase normalised to peak from (E). Statistics: \**p*<0.05, \*\**p*<0.01, \*\*\**p*<0.001, two-way repeated measures ANOVA (C,D), and one-way ANOVA with Sidak multiple comparisons (E). Nonlinear regression with extra-sum-of-squares F test; *τ* = decay time constant (F). Error bars indicate SEM.

To further characterise and validate the role of ENT1 and astrocytes in striatal adenosine signalling, we explored their impact on the dynamics of extracellular adenosine transients evoked electrically by trains of electrical stimulation (100 pulses, 50 Hz) (**Figure 5E**). Evoked increases in extracellular adenosine concentrations exhibited relatively extended rise times (~30 sec) and clearance times as reported previously in cultured hippocampal neurons and acute hippocampal and medial prefrontal cortex mouse brain slices (Wu et al., 2020), and surprisingly release was activated via a mechanism that was not prevented by Na_v_ blocker TTX (1 μM). (**Figure 5E**, F_(3,29)_ = 13.60, *p* < 0.0001, one-way ANOVA; TTX vs control: *p* > 0.99, Sidak’s multiple comparisons). In slices pre-treated with the ENT1 inhibitor NBTI (10 μM, for 45 – 60 min), the peak of evoked adenosine levels was attenuated (**Figure 5E**, F_(3,29)_ = 13.60, *p* < 0.0001, one-way ANOVA; NBTI vs control: *p* = 0.001, Sidak’s multiple comparisons), but the clearance time constant was extended (**Figure 5F**, F = 8543, *p* < 0.0001, extra-sum-of-squares F test; control: τ = 108.4, NBTI: τ = 564.5), indicating reduced release and uptake, and seen previously in cultured hippocampal neurons (Wu et al., 2020). In slices pre-treated with the gliotoxin fluorocitrate (100 μM, for 45 – 60 min), both the peak level of adenosine and the time constant for clearance were elevated (**Figure 5E**, F_(3,29)_ = 13.60, *p* < 0.0001, one-way ANOVA; FC vs control: *p* = 0.049, Sidak’s multiple comparisons; **Figure 5F**, F = 8543, *p* < 0.0001, extra-sum-of-squares F test; control: τ = 108.4, FC: τ = 217.0), indicating that ENT1 on astrocytes support adenosine uptake

### EtOH increases tonic A_1_R-mediated inhibition of DA release

Our data indicate that astrocytic ENT1 function regulates tonic A_1_R-mediated inhibition of DA output in NAcC. Acute exposure to ethanol is documented to increase extracellular adenosine levels in many brain nuclei, including striatum, via inhibition of adenosine uptake by ENT1 (Nagy et al., 1989, 1990; Choi et al., 2004). Given that ethanol also attenuates evoked DA release in NAc (Yorgason et al., 2014, 2015; Karkhanis et al., 2015; Rose et al., 2016), we tested whether acute ethanol exposure might increase tonic A_1_R-mediated inhibition of DA release via impaired ENT1 function. We pre-treated slices with ethanol (2–3 hrs), at a concentration (50 mM) that correlates to a blood alcohol concentration of 230 mg/dl in humans and is consistent with what late-stage alcoholics achieve (Brick and Erickson, 2009). We first confirmed that pre-treating slices with ethanol using this paradigm reduced [DA]_o_ evoked by single and five pulses (50 Hz) of electrical stimulation in NAcC (**Figure 6A**, 1p: t_(12)_ = 2.74, *p* = 0.018; 5p: t_(12)_ = 2.67, *p* = 0.020; unpaired t tests). Next, we found that the effect of A_1_R antagonism with CPT (10 μM) on [DA]_o_ evoked by single electrical pulses was elevated compared to control slices (**Figure 6B**, F_(1, 12)_ = 18.04, *p* = 0.0011, two-way RM ANOVA: main effect of drug). A_1_ receptor antagonism with CPT decreased the ratio of [DA]_o_ evoked by 5p:1p (50 Hz) (**Figure 6C**, F_(1, 12)_ = 68.51, *p* < 0.0001, two-way RM ANOVA: main effect of drug), and there was a significant interaction between ethanol and CPT (**Figure 6C**, F_(1, 12)_ = 9.36, *p* = 0.0099, two-way RM ANOVA: ethanol x CPT interaction), which was borne out by a more pronounced decrease after ethanol than in control slices (**Figure 6D**, t_(12)_ = 3.16, *p* = 0.0082, unpaired t test). To test directly whether ethanol in this paradigm impaired adenosine uptake by ENT1, we imaged tonic extracellular adenosine levels with the GRAB-Ado sensor during application of ENT1 inhibitor NBTI (10 μM). NBTI increased tonic extracellular adenosine levels in NAcC to a significantly lesser degree in slices preincubated with ethanol than controls (**Figure 6E**, F_(1,17)_ = 11.30, *p* = 0.0037, two-way RM ANOVA: main effect of drug; t_(17)_ = 2.60, *p* = 0.019, unpaired t test). These data indicate that tonic A_1_R-mediated inhibition of DA axons by adenosine in the NAcC is elevated by acute ethanol exposure, paralleled by an underlying attenuated uptake of adenosine uptake by ENT1.

**Figure 6.**
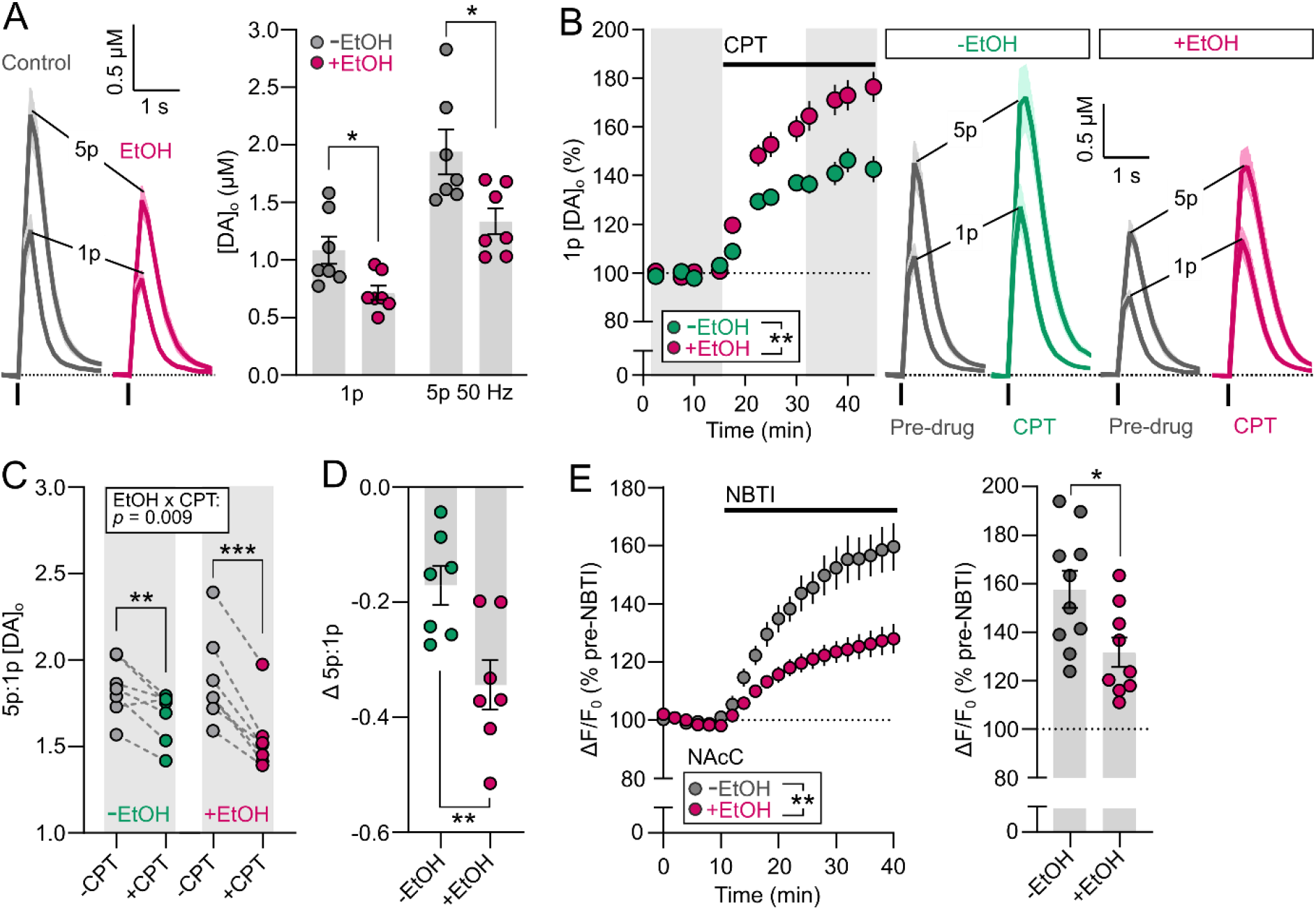
EtOH enhances tonic adenosine A_1_R-mediated inhibition of DA release. **A**, Mean [DA]_o_ transients (left) and corresponding mean peak [DA]_o_ (right) evoked by a single electrical pulse (1p) or five pulses (5p) at 50 Hz in NAcC in the absence (grey; *n* = 7 experiments/4 mice) or presence of 50 mM EtOH (pink; *n* = 7 experiments/4 mice). **B**, Summary of mean peak [DA]_o_ before (grey) and after application of A_1_R antagonist CPT (10 μM) normalised to pre-drug baseline (dotted line) (left) and right, corresponding mean [DA]_o_ transients evoked by a single electrical pulse (1p) and five pulses (5p) at 50 Hz in NAcC in the absence (green, *n* = 7 experiments/4 mice) or presence of 50 mM EtOH (pink; *n* = 7 experiments/4 mice). **C-D**, Ratio of peak [DA]_o_ evoked by 5p:1p at 50 Hz (C) and Δ 5p/1p (D) before (grey) and after application of CPT (10 μM) in absence (left, green) or presence of 50 mM EtOH (right, pink). Data summarised from (B). **E**, Summary (left) and mean peak (right) of unstimulated ΔF/F_0_ GRAB-Ado signal in NAcC normalised to pre-drug baseline (dotted line) before and after application of ENT1 inhibitor NBTI (10 μM) in the absence (grey; *n* = 10 experiments/5 mice) or presence of 50 mM EtOH (pink; *n* = 9 experiments/5 mice). Statistics: *p*<0.05, \*\**p*<0.01, \*\*\**p*<0.001, unpaired *t* test (A,D), two-way repeated measures ANOVA (B,C,E) with Sidak multiple comparisons (C). Error bars indicate SEM.

## DISCUSSION

Here, we reveal that DA release in NAcC is under a tonic inhibition by ambient adenosine levels acting at A_1_Rs, and that ENT1, located at least in part on striatal astrocytes, governs the level of this tonic inhibition. Moreover, we reveal that ethanol promotes A_1_R-inhibition of DA release, through elevating adenosine levels by diminishing adenosine uptake via ENT1. These data support the emerging concept that astrocytes play important roles in setting the level of striatal DA output, in health and disease.

### Direct versus indirect actions of A_1_Rs

Our data provide functional evidence for direct regulation of DA release by A_1_Rs. Immunocytochemical studies in rat striatal synaptosomes indicate that dopaminergic axons can contain A_1_Rs (Borycz et al., 2007), but direct immunocytochemical evidence *in situ* is currently lacking. We excluded indirect actions of A_1_Rs via key candidate pathways, namely cholinergic inputs that act via nAChRs and GABAergic networks that act via GABAA/GABAB receptors on DA axons. Previous reports have excluded effects of glutamatergic modulation (Borycz et al., 2007; O’Connor and O’Neill, 2008). Moreover, we found that A_1_R agonists inhibited DA release evoked by even single short optogenetic stimulation of ChR2-expressing DA axons, which should not co-activate other striatal neurons, suggesting that the location(s) for A_1_Rs that regulate DA are either a currently undisclosed tonically active cell type with currently unknown actions on DA or, more parsimoniously, DA axons themselves.

### Tonic inhibition

A_1_R antagonists enhanced DA release evoked by single short optogenetic stimulation, suggesting that A_1_R activation does not require stimulation of any other network, and is tonically activated by an ambient striatal adenosine tone. An adenosine tone at striatal A_1_Rs has previously been described on glutamate inputs in NAcC (Brundege and Williams, 2002; Choi et al., 2004). Extracellular adenosine concentrations in the brain have been suggested to be in the range of 25–250 nM under basal conditions, which are sufficient to activate high-affinity A_1_Rs (Dunwiddie and Masino, 2001). In further support for a resting adenosine tone in NAcC, we could detect extracellular adenosine with the GRAB-Ado sensor, following application of an ENT1 inhibitor, in the absence of any striatal stimulation. The source of adenosine is still undetermined, but can arise from the catabolism of ATP from neuronal and non-neuronal sources (Latini and Pedata, 2008), including astrocytes (Corkrum et al., 2020).

### A_1_Rs modify DA signal contrast for firing frequency

We found that A_1_R activation not only limits the overall amplitude of DA output, but promotes the contrast in DA signals released by different firing rates and pulse numbers, and vice versa, A_1_R antagonists reduce this contrast. The strength of A_1_R-mediated inhibition of DA release varied with adenosine concentration, resulting from ENT1 inhibition or overexpression, and furthermore, was greater for lower frequencies of activation of DA axons. These data indicate that an adenosine tone preferentially limits DA output during the lower frequencies of DA neuron activity that represent tonic activity, while leaving relatively intact the DA release evoked by the higher frequencies associated with phasic activity in DA neurons. The preferential inhibition of DA release by low frequencies of activity parallels the effects of GABA input to DA axons (Lopes et al., 2019; Roberts et al., 2020; Roberts et al., 2021) and could indicate a preferential influence of striatal adenosine on DA functions that are proposed to be mediated by low frequencies of activity e.g. the ongoing monitoring of reward value and its changes (Wang et al., 2021). We identified an underlying change in the dynamic short-term plasticity (STP) of DA output that corresponds to this change to frequency filtering, that again parallels the effect of GABA on STP in DA release (Lopes et al., 2019; Roberts et al., 2020). Elsewhere in the brain, A_1_R activation, including through ambient adenosine, can modulate STP of glutamate transmission through a presynaptic mechanism (Perrier et al., 2019; Qi et al., 2017; Yang et al., 2007; Moore et al., 2003; Wong et al., 2006). A_1_Rs are G_i/o_ coupled, and so their activation causes inhibition of adenylyl cyclase, activation of potassium channels and inactivation of voltage-gated calcium channels (VGCCs) (Haas and Selbach, 2000). We have recently revealed that mechanisms that determine axonal excitability, particularly potassium-dependent processes, strongly gate STP of DA release (Condon et al., 2019) while calcium largely gates amplitude. We therefore hypothesise that A_1_Rs on striatal DA axons likely inhibit the overall amplitude of DA release via reduced VGCC activity while simultaneously gating the STP of DA release by modifying axonal excitability via potassium-dependent conductances. Future studies are needed to establish these potential mechanisms.

### ENT1 and astrocytes are key regulators of tonic A_1_R-inhibition of DA release

We established that ENT1 in NAcC limits ambient adenosine levels and tonic A_1_R-inhibition of DA axons, and thereby indirectly facilitates DA release. ENT1 usually facilitates adenosine uptake from the extracellular milieu, but can reverse to release adenosine (Parkinson et al., 2011), and ENT1 has been shown to facilitate adenosine release evoked by electrical stimulation in cultured hippocampal neurons (Wu et al., 2020). Intriguingly, we found that adenosine release driven by electrical stimulation was TTX-insensitive and sensitive to ENT1 inhibition. Together, our data show that ambient adenosine in NAcC is limited by uptake via ENT1, but can be released via ENT1 reversal in some conditions. There is an additional ENT in striatum, ENT2 (Jennings et al., 2001; Anderson et al., 2002), whose roles we did not explore. ENT1 is thought to be responsible for the majority of adenosine transport and, subsequently, the key regulator of extracellular adenosine levels across the brain (Young et al., 2008).

We found a major role for ENT1 located to striatal astrocytes, although we did not test or exclude roles for ENT1 located on neurons. The glial metabolic poison fluorocitrate limited the effects of ENT1 inhibitors on adenosine tone, while overexpressing ENT1 in astrocytes boosted evoked DA release by limiting tonic A_1_R-inhibition. This strategy for viral overexpression of ENT1 in striatal astrocytes has previously been shown to increase ENT1 expression by ~30%, but also increases striatal GFAP expression and modifies astrocyte morphology (Hong et al., 2020). ENT1 expression has also been shown to regulate astrocyte-specific excitatory amino acid transporter 2 (EAAT2) and aquaporin-4 expression in NAcC (Wu et al., 2010; Lee et al., 2013). These additional astrocyte-specific modifications might also contribute to boosted DA signalling in NAcC, beyond mechanisms involving tonic inhibition by adenosine tone. Regardless, astrocytic ENT1 in striatum has previously been reported to play key roles in reward-seeking behaviours (Hong et al., 2020; Kang et al., 2020), which our data suggest could be mediated by underlying changes to DA signalling.

The role for ENT1 on astrocytes in gating DA output parallels our recent finding that GABA transporters (GAT1 and GAT3) on astrocytes in dorsal striatum set the level of tonic GABAergic inhibition of DA release (Roberts et al., 2020). Furthermore, EAAT2, enriched on astrocytes, limits glutamate-mediated inhibition of DA release (Zhang and Sulzer, 2003), and thus, our collective findings point to astrocytic transporters across neurotransmitter categories as regulators of DA release.

### Dysregulation of A_1_R-inhibition of DA release by ethanol

To probe the wider potential significance of the regulation of DA in NAcC by adenosine tone and ENT1, we explored whether A_1_R inhibition of DA release was modified by ethanol. Ethanol has been shown to increase extracellular adenosine levels by impairing adenosine uptake via ENT1 after acute exposure (Nagy et al., 1989, 1990; Choi et al., 2004), and separately, to reduce DA release following acute application (Yorgason et al., 2014, 2015) or following chronic intermittent exposure in mice (Karkhanis et al., 2015; Rose et al., 2016). Here we bridge these different findings by revealing that ethanol can reduce DA output via a boosted tonic A_1_R-inhibition. While pre-treating slices with 50 mM ethanol to understand disruption to cellular and circuit function has limitations in informing effects on behaviour, we speculate that this mechanism could help to explain how striatal A_1_Rs contribute to the ataxic and hypnotic effects of ethanol (Meng and Dar, 1995; Phan et al., 1997; Dar, 2001), as well as how acute ethanol exposure attenuates concurrent GABA co-release from DA axons in dorsal striatum (Kim et al., 2015).

Chronic ethanol exposure is thought to evoke an adaptive response, resulting in decreased ENT1 expression, and therefore a reduced ability for ethanol to increase extracellular adenosine (Nagy et al., 1989). Indeed, rats permitted daily access to ethanol for 8 weeks exhibit downregulated ENT1 gene expression in NAc (Bell et al., 2009). ENT1-null mice exhibit reduced hypnotic and ataxic responses to ethanol, increased ethanol consumption, and decreased adenosine tone in NAc, while viral-mediated rescue of ENT1 expression in NAc reduces ethanol consumption (Choi et al., 2004; Jia et al., 2020). Furthermore, growing evidence implicates striatal adenosine signalling in the neurobiological adaptations of other drugs of abuse (Bachtell, 2017; Ballesteros-Yáñez et al., 2018). Repeated cocaine administration enhances adenosine uptake, reduces adenosine tone, and reduces plasma membrane A_1_R expression in NAcC (Manzoni et al., 1998; Toda et al., 2003), while striatal μ-opioid receptor activation by opioids results in reduced striatal adenosine tone in dorsomedial striatum (Adhikary and Birdsong, 2021). Dysregulated tonic A_1_R-inhibition of glutamate afferents in NAc is thought to be a key circuit adaptation underlying ethanol abuse and other drugs of abuse (Choi et al., 2004; Chen et al., 2010; Wu et al., 2010; Nam et al., 2011); however, given the role adenosine tone plays in setting the level and activity-dependence of DA output we describe here, we speculate that dysregulated tonic A_1_R-inhibition of DA release in NAcC might also be an important circuit adaptation underlying drug abuse.

In conclusion, we show here that A_1_Rs can tonically inhibit DA output though an ambient adenosine tone, and that A_1_Rs additionally regulate the activity-sensitivity of DA release, preferentially impacting on release by low frequencies. Furthermore, we find that tonic A_1_R-inhibition of DA release is regulated by ENT1 and astrocytes, and dysregulated by ethanol. These data provide a further mechanism through which ethanol modulates striatal DA function, and corroborate emerging data highlighting astrocytic transporters as important regulators of striatal function.

## Acknowledgements

This work was supported by grants from the UK Medical Research Council (MR/V013599/1 to S.J.C. and B.M.R.) and the University of Oxford John Fell Fund (0008310 to B.M.R.), a Junior Research Fellowship to B.M.R from St John’s College, Oxford, and a BBSRC Doctoral Training Grant to J.A.L.

